# Multi-level Prediction and Landscape Characterization of Human *γδ* T Cells Across Single-Cell and Spatial Transcriptomic Atlases

**DOI:** 10.64898/2026.11.24.769075

**Authors:** Ka Ning Wen, Yuanhua Huang, Sunghee Park

**Affiliations:** School of Biomedical Sciences, The University of Hong Kong, Hong Kong, China; School of Computing and Data Science, The University of Hong Kong, Hong Kong, China; Department of Biomedical Systems, Soongsil University, Seoul, Republic of Korea

## Abstract

Human *γδ* T cells contribute to tissue immunity and tumour responses, but their identification in single-cell transcriptomic data is limited by incomplete T-cell receptor (TCR) recovery and transcriptional overlap with *αβ* T cells and NK cells. Here we present gdTProfiler, a transcriptome-based framework for *γδ* T-cell identification, V*δ*1/V*δ*2 subtype prediction and cell-state characterization. A joint classifier distinguished *γδ* T, *αβ* T and NK cells with an average precision of 0.99 in donor-held-out and external datasets, and predicted V*δ*1/V*δ*2 subtype from non-TCR gene expression. In NK-annotated atlas populations, the model recovered all 25 cells with linked TRG/TRD sequences as *γδ* T cells, and these cells showed higher cytotoxic programme usage. Consensus non-negative matrix factorization identified eight transcriptional programmes that captured stimulus-associated variation and organized a cross-dataset atlas of 182,638 *γδ* T-cell records. Applied to colorectal cancer spatial transcriptomics data, gdTProfiler identified candidate *γδ* T-cell-associated regions for follow-up. gdTProfiler provides an integrated framework and reusable reference for studying *γδ* T-cell identity and heterogeneity across single-cell and spatial datasets.

---

Human *γδ* T cells contribute to tissue immune surveillance and to responses against infection and malignancy.^1–3^ Their TCR subtype composition and transcriptional states vary across tissues and throughout life. ^4^ V*δ*-chain usage distinguishes the major human *γδ* T-cell subsets, including V*δ*1^+^ and V*δ*2^+^ cells, but subtype alone does not define cellular states: cells within the same subtype can occupy distinct functional states characterized by differences in differentiation, activation and cytotoxic programmes.^5,6^ Describing this diversity therefore requires *γδ* T-cell identity, TCR subtype and transcriptional state to be resolved together.

Single-cell transcriptomics can capture this heterogeneity across tissues, but *γδ* T cells are difficult to annotate when TCR information is incomplete. In mixed peripheral blood mononuclear cells, initial low-resolution clustering did not resolve a distinct *γδ* T-cell cluster, and flow-cytometry-purified cells were needed to define their transcriptional signatures. ^5^ Paired TCR sequencing provides direct evidence of lineage and subtype, but such information is available for only a subset of cells and datasets. Effector programmes shared with *αβ* T cells and natural killer (NK) cells further limit the extent to which transcriptional similarity alone can substitute for TCR-defined identity.^5,7^ As a result, *γδ* T cells can be misclassified as NK cells or remain unresolved from other low-abundance lymphocyte populations, motivating a supervised approach for *γδ* T-cell identification.

Existing annotation strategies address different aspects of this problem. Marker- and module-based approaches, including the modular TCR-expression score of Song et al. ^8^ and gating tools such as scGate, ^9^ rely on a small number of TCR-associated and lineage genes and therefore depend on their detection in each cell. Reference classifiers such as CellTypist ^10^ and label-transfer methods such as scANVI and Azimuth ^11,12^ use broader expression profiles, but they learn from transcriptionally defined annotations rather than TCR-defined labels and are not designed to separate V*δ*1 from V*δ*2 cells. A classifier trained directly on TCR-supported labels, evaluated across donors and cohorts, and able to distinguish *γδ* T cells from both *αβ* T cells and NK cells would complement these approaches.

Comparing *γδ* T-cell states across datasets also requires a shared representation of transcriptional variation. Gene-expression programmes complement discrete labels because multiple transcriptional activities can coexist within a single cell, and consensus non-negative matrix factorization (cNMF) estimates both the programmes and their contributions to each transcriptome.^13^ A fixed programme reference would allow states to be compared across cohorts and enable new cells to be mapped into a common space. Spatial transcriptomics could then extend this framework to their spatial context within tissues, provided that transfer across measurement platforms is carefully evaluated.

Here we present gdTProfiler, a framework for multi-level prediction and landscape characterization of human *γδ* T cells. A joint model trained on TCR-supported labels separates *γδ* T, *αβ* T and NK cells and predicts V*δ*1/V*δ*2 subtype from non-TCR expression. We evaluate the model using donor-held-out testing and independent external cohorts, and apply it to NK-annotated populations in two cell atlases, where linked TCR sequences confirm a subset of the predicted *γδ* T cells. We derive eight cNMF programmes from adult *γδ* T cells, show that they capture stimulus-associated variation, and use them to assemble a cross-dataset cell-state atlas for mapping new transcriptomes. Finally, we apply gdTProfiler to spatial transcriptomes from colorectal cancer to localize candidate *γδ* T cells in colorectal cancer tissue.

## Results

### The workflow for gdTProfiler

We developed gdTProfiler to link *γδ* T-cell identification and TCR-subtype prediction with a shared description of transcriptional state (Fig. 1). An identity head classifies cells as *αβ* T, *γδ* T or NK cells, and a parallel subtype head predicts V*δ*1 versus V*δ*2 from non-TCR gene expression. A separate cell-state programme reference describes continuous expression variation and supports atlas mapping. The model was trained on paired-TCR and TRD-only Song cells and evaluated on paired-TCR reference cells (Fig. 1a; Extended Data Fig. 1). We assessed performance in donor-held-out and external cohorts, and NK specificity in held-out patients.

**Figure 1:**
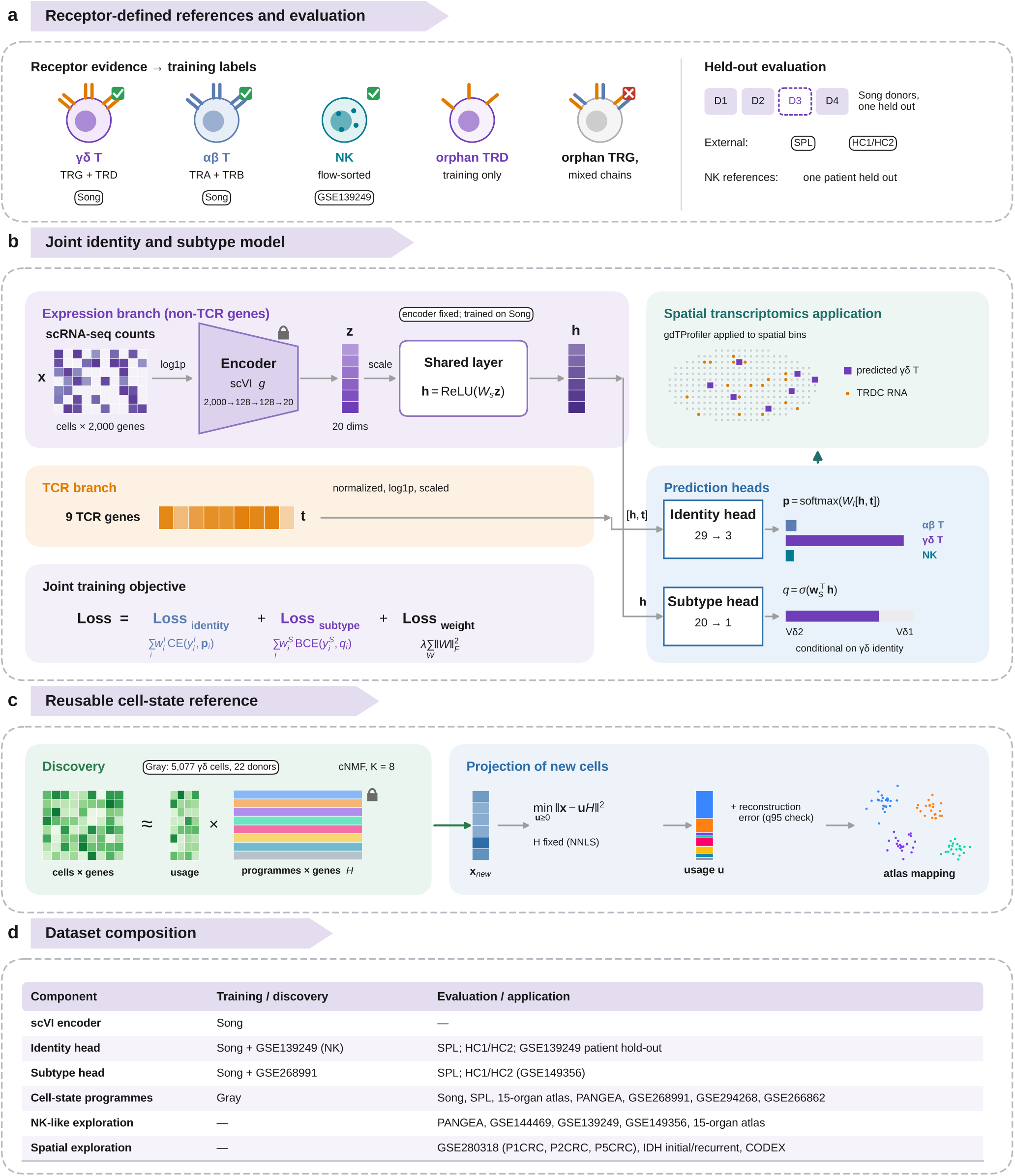
gdTProfiler: joint identity and subtype prediction with a reusable cell-state reference. **a,** TCR evidence defines training labels and held-out evaluation. Paired TRG+TRD without TRA/TRB and paired TRA+TRB without TRG/TRD define *γδ* and *αβ* references, respectively, and flow-sorted GSE139249 cells provide NK references. Song cells with TRD-only evidence are used for training only and are excluded from paired-reference testing; TRG-only and mixed-chain records are not given definite *γδ* labels. Song evaluation holds out one donor at a time, SPL and HC1/HC2 are external references never used for training, and NK references are assessed by holding out one patient at a time. **b,** Non-TCR expression enters a fixed scVI encoder trained on Song; the 20-dimensional representation is standardized and passed through a trainable shared 20 *→* 20 ReLU layer. Nine TCR features are concatenated with the shared representation only for the identity head (29 *→* 3, softmax over *αβ* T, *γδ* T and NK). The subtype head (20 *→* 1, sigmoid) uses the shared representation alone and estimates V*δ*1 versus V*δ*2 conditional on *γδ* identity. The shared layer and both heads are trained jointly with a weighted identity cross-entropy, a weighted subtype binary cross-entropy over labelled cells and an L2 weight penalty (Methods). The fixed model is applied without refitting to atlas cells and spatial transcriptomic bins, providing the NK-labelled atlas predictions in Fig. 3 and the spatial predictions in Fig. 6. **c,** Eight programmes learned by consensus NMF from 5,077 Gray *γδ* T cells across 22 donors are kept fixed. New cells are projected by non-negative least squares with the programme spectra held fixed, giving programme usage for exploratory atlas mapping and a reconstruction error that assesses applicability. **d,** Training or discovery datasets and evaluation or application datasets for each component. GSE268991 does not train the encoder or programme basis. Conditional subtype performance does not establish end-to-end identity-gated recovery. Expression matrices, probability bars, spatial bins and embeddings are schematic.

### Joint prediction identifies *γδ* T cells across donors and cohorts

To evaluate generalization across donors, we performed leave-one-donor-out validation in the Song cohort. ^8^ The joint model identified *γδ* T cells with an average precision (AP) of 0.99 (Fig. 2a,b). Performance was maintained in an independent spleen dataset (SPL) from the Cross-tissue Immune Cell Atlas (AP = 0.99; Fig. 2d), with 91% recall and 98% precision at a probability threshold of 0.5 (Fig. 2g,h).^10^ Because NK cells share cytotoxic and innate-like transcriptional programmes with *γδ* T cells, we further assessed specificity in flow-sorted NK cells using patient-held-out models: 843 of 848 cells were classified as NK and none as *γδ* T cells. ^14^ The model also identified *γδ* T cells lacking detectable *TRDC* transcripts, and its performance was stable across random seeds (Extended Data Fig. 2).

**Figure 2:**
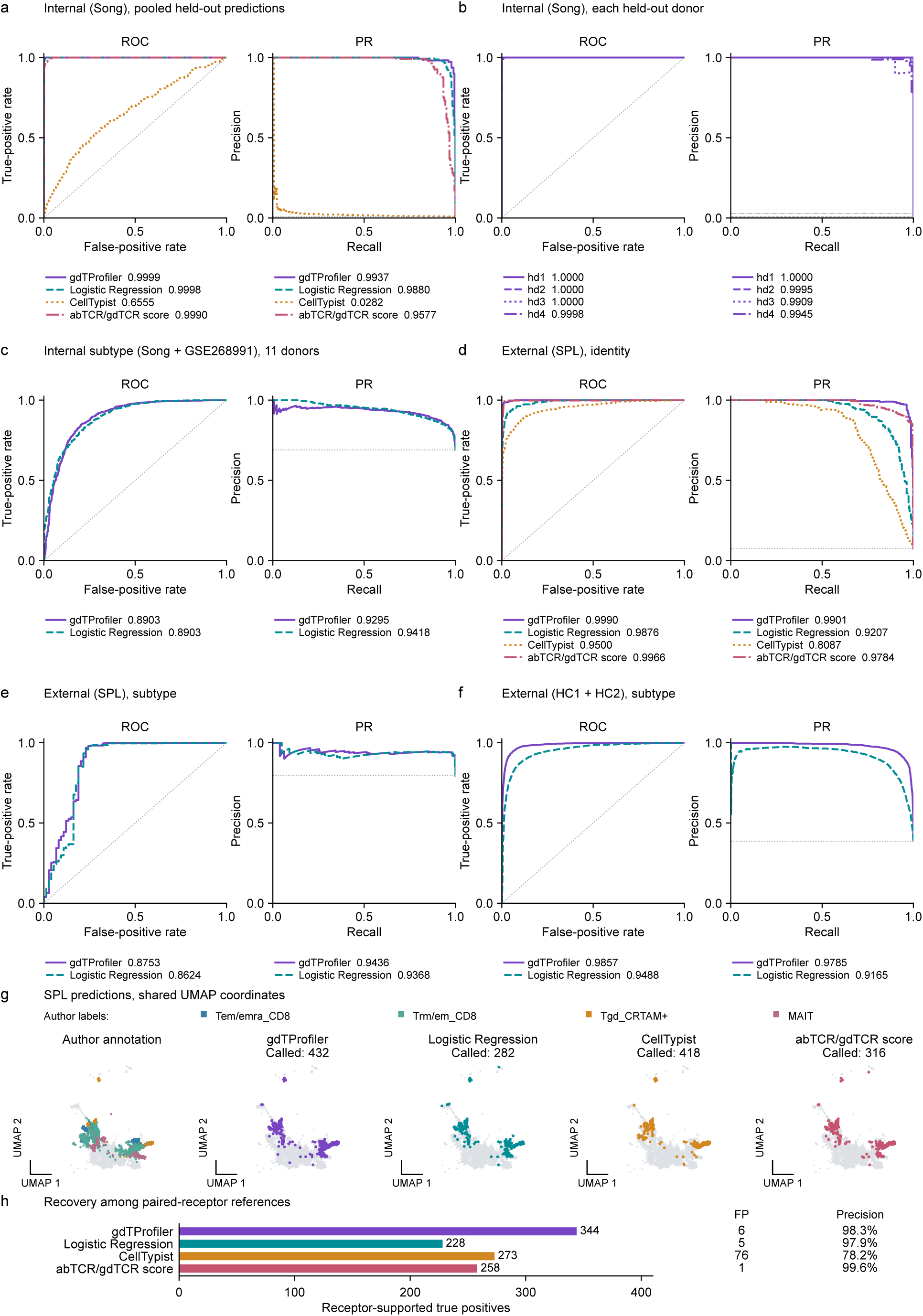
gdTProfiler identity and conditional subtype prediction across donors and cohorts. **a,b,** Song identity ROC and precision–recall curves, pooled across four donor-held-out fits (a) and shown separately for each donor for gdTProfiler (b). Each fold’s encoder, scalers and supervised warm start excluded the held-out donor. The 20,254 paired references comprise *γδ/αβ* counts of 10/6,369 (hd1), 65/6,355 (hd2), 10/3,965 (hd3) and 97/3,383 (hd4). Line styles distinguish donors in b; hd1 and hd3 each have only ten positive references. **c,** Conditional subtype internal validation across 11 Song+GSE268991 donors (1,511 V*δ*1 and 677 V*δ*2 paired references), excluding every held-out donor’s cells from supervised fitting and warm starts. **d,** SPL identity assessment (378 *γδ*, 4,668 *αβ*; five donors). **e,f,** Conditional subtype assessment in SPL (288 V*δ*1, 74 V*δ*2) and HC1+HC2 (3,707 V*δ*1, 5,881 V*δ*2), respectively. Identity uses unconditional *p*(*γδ*); subtype uses *q*(V*δ*1) within TCR-defined references irrespective of identity score. Legends give AUROC or AP; grey reference lines indicate chance ranking or prevalence, separately for each donor in b. HC identity recovery is shown in Extended Data Fig. 2. **g,** Author annotations and predictions on all 5,198 SPL cells in the same UMAP coordinates. Original Tgd_CRTAM+, Tem/emra_CD8, Trm/em_CD8 and MAIT annotations are gold, blue, green and rose; other annotations are grey. Author labels were matched by cell identifier and describe transcriptional annotations rather than TCR-defined identity. **h,** True positives, false positives (FP) and precision among paired SPL references. gdTProfiler and Logistic Regression use a 0.5 probability threshold; CellTypist uses its native label. The abTCR/gdTCR score adapts Song gene panels and Seurat AddModuleScore using the minimum margin to two source-training-derived gates. Internal gates exclude the held-out donor; SPL gates use source Song cells without SPL labels. Logistic Regression includes TCR expression for identity and excludes it for subtype. Comparators use matched cells but differ in supervision. One seed (17); SPL and HC1/HC2 were excluded from supervised fitting but inspected during development and constitute retrospective external assessments.

We next evaluated V*δ*1/V*δ*2 subtype prediction from non-TCR gene expression. Under leave-one-donor-out validation across 11 donors, gdTProfiler performed comparably to a logistic regression baseline (AUROC = 0.89 for both; Fig. 2c). In external spleen and peripheral blood cohorts (healthy donors HC1 and HC2), gdTProfiler achieved AP values of 0.94 and 0.98, respectively, compared with 0.94 and 0.92 for logistic regression (Fig. 2e,f).^10,15^ In the blood cohorts, which contain only *γδ* T cells, the model recovered more than 98% of cells as *γδ* T cells (Extended Data Fig. 2). These results indicate that *γδ* T-cell identity and subtype can be inferred from the transcriptome alone across donors and tissues.

### gdTProfiler detects NK-like *γδ* T cells within NK-labelled populations

*γδ* T cells share cytotoxic and innate-like transcriptional features with NK cells. ^5,7^ We therefore tested whether *γδ* T cells were present among NK-annotated cells in two public atlases, the 15-organ atlas of He et al. ^16^ and the PANGEA atlas of Kim et al. ^17^ The model identified 83 of 2,586 NK/T-labelled cells in the 15-organ atlas and 118 of 13,677 NK-labelled cells in PANGEA as *γδ* T-cell candidates (Fig. 3a–f). Comparison with the original annotations in the same embedding highlighted cells for annotation reassessment (Extended Data Fig. 3).

**Figure 3:**
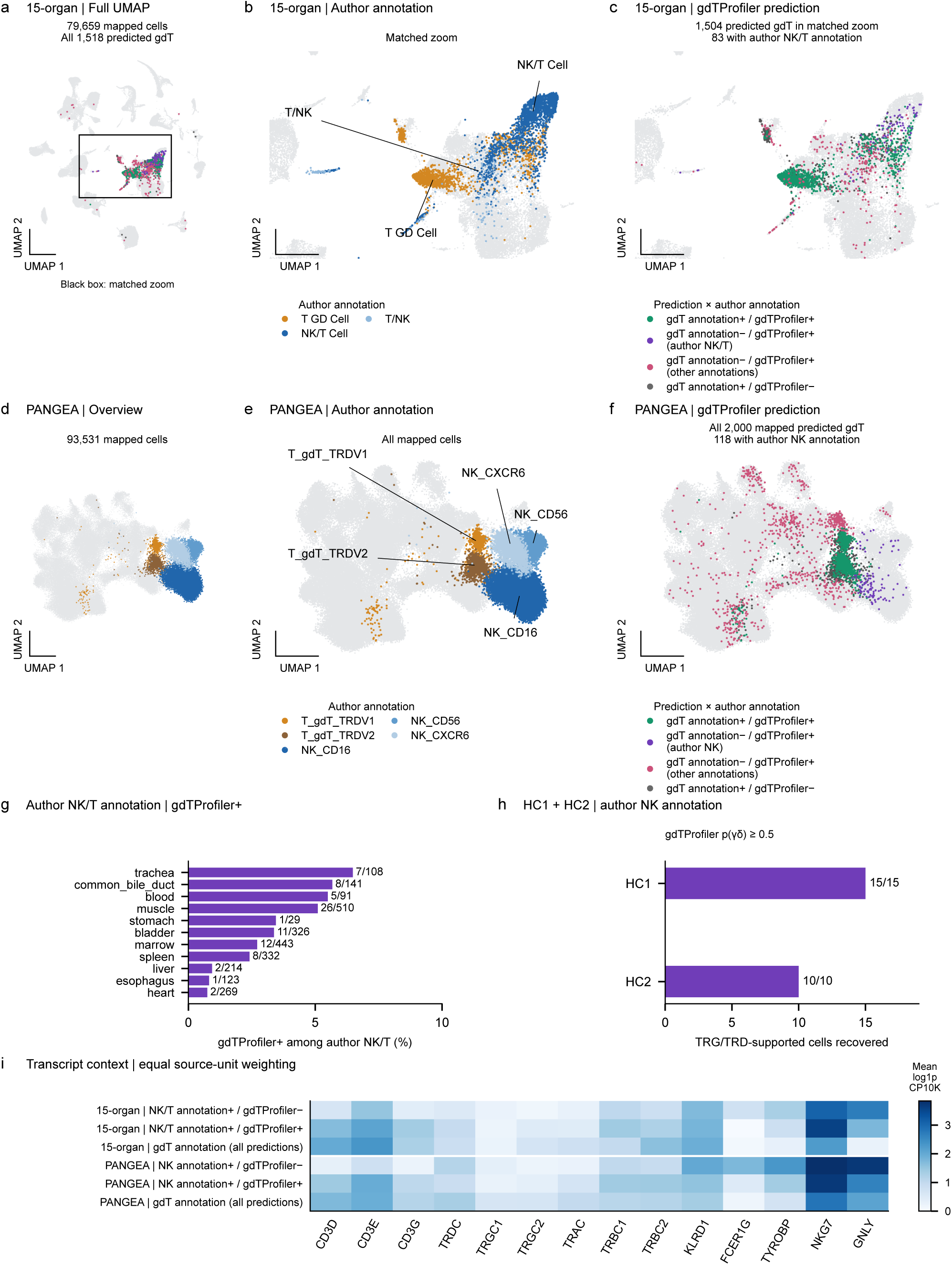
gdTProfiler TCR-supported recovery within NK-labelled atlas populations. Predictions use the same three-class gdTProfiler model as Fig. 2. **a,** Complete 15-organ prediction UMAP (79,659 mapped records), showing all 1,518 mapped predicted-positive cells. The black rectangle marks the matched zoom in b,c. **b,c,** Enlarged author-annotation and gdTProfiler-prediction maps of the T GD Cell, NK/T Cell and T/NK neighbourhood, with identical coordinates, limits, dimensions and aspect ratio. The zoom contains 1,504 predicted-positive records, including 83 originally NK/T-labelled records; the remaining 14 mapped predictions stay visible in a. The window illustrates existing author-labelled geometry and does not change quantitative denominators. **d–f,** Complete PANGEA overview (93,531 mapped records), author annotations and predictions. Separate legends and palettes distinguish annotation from prediction. Author maps use gold/brown for published *γδ* labels and blue shades for NK/T or NK labels. In 15-organ, the NK/T group comprises the original NK/T Cell and T/NK labels, which remain distinct on the author map. Prediction maps show mutually exclusive groups: green, gdT annotation+/gdTProfiler+; purple, gdT annotation-negative/gdTProfiler+ with author NK/T or NK annotations; rose, gdT annotation-negative/gdTProfiler+ with other author annotations; dark grey, gdT annotation+/gdTProfiler-negative. Remaining cells are light grey. The three positive groups sum to the displayed prediction count. TRG/TRD-support symbols are omitted from these maps; TCR-supported recovery is quantified in panel h. Full prediction panels a and f retain all 1,518 mapped 15-organ and 2,000 mapped PANGEA predicted-positive records, respectively, without subsampling. Records lacking coordinates remain in source tables and quantitative summaries; no positions are imputed. **g,** gdTProfiler-positive fractions within author NK/T annotations by organ, with numerator/denominator labels. **h,** Recovery of originally NK-labelled HC1/HC2 cells with matched processed TRG/TRD and CDR3 evidence: 15/15 and 10/10, respectively. These PANGEA-processed cells come from the same HC1/HC2 donors assessed using the complete GEO matrices in Fig. 2; they are not an additional cohort. **i,** Marker expression for author NK/T or NK annotations stratified by gdTProfiler prediction, and for all author gdT annotations irrespective of prediction. Expression is normalized per complete library, averaged within source units and then equally across units. Fifteen-organ units are organs, not donors; PANGEA units are study/sample combinations. Calls use unconditional softmax *p*(*γδ*) *≥* 0.5. PANGEA includes GSE139249, which supplied NK training references; full-atlas application is therefore not wholly external validation. Original annotations and threshold/TCR-conflict checks are provided in Extended Data Figs. 3 and 4.

To validate these predictions, we examined cells with linked TCR sequences. In the PANGEA-processed subset of the HC1 and HC2 donors, all 25 NK-labelled cells with matched TRG, TRD and CDR3 sequences were predicted as *γδ* T cells (Fig. 3h). Candidates co-expressed T-cell and NK-associated markers (Fig. 3i). Candidates with unavailable or discordant TCR evidence, including cells with paired *αβ* TCRs, are reported separately (Extended Data Fig. 4). These results indicate that NK-annotated populations in public atlases include *γδ* T cells, and that gdTProfiler can prioritize such cells for re-annotation.

### The cell state model detects stimulus-associated transcriptional variation

To characterize *γδ* T-cell state independently of TCR subtype, we derived eight consensus programmes from the adult *γδ* T-cell reference of Gray et al.,^4^ comprising 5,077 cells from 22 donors (Fig. 4a; Extended Data Fig. 5). Based on their leading genes, these included a cytotoxic programme (P2; *NKG7*, *GZMH*, *PRF1*), an immediate-early response programme (P5; *CD69*, *FOS*, *NFKBIA*), a heat-shock programme (P7; *HSPA1A*, *DNAJB1*) and a cell-cycle programme (P9; *CDC20*, *PLK1*), consistent with hallmark gene-set overlap (Fig. 4b). P3 was led by *TMIGD2*, *LDLRAD4* and *ITGA1*.

**Figure 4:**
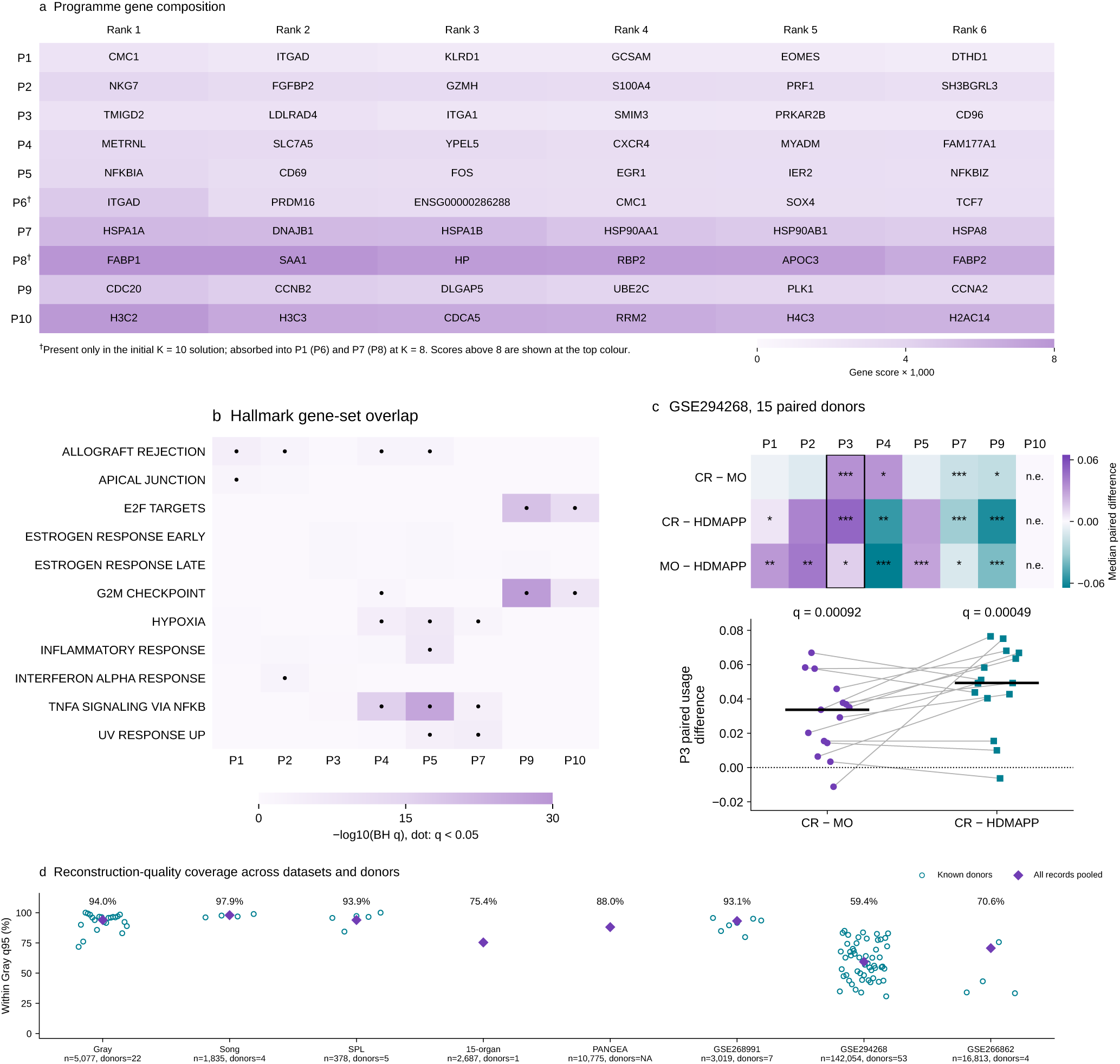
Cell-state programmes, stimulus-associated variation and projection applicability. **a,** Six highest-scoring genes in each Gray *K* = 8 programme; colour denotes gene score multiplied by 1,000. Labels follow the matched programmes of an initial *K* = 10 solution; *K* = 10 programmes P6 and P8 were absorbed into P1 and P7, respectively (Methods); rows P6 and P8 (*†*) show the six highest-scoring genes of these *K* = 10 programmes, and scores above 8 are shown at the top colour. **b,** Hallmark overlap using the top 50 genes per programme. Colour denotes *−* log_10_ adjusted *P* ; dots indicate adjusted *P <* 0.05, with Benjamini–Hochberg correction across 800 tests. Gene overlap supports descriptive interpretation, not functional validation. **c,** Programme usage after cockroach extract (CR), mouse extract (MO) and the phosphoantigen HDMAPP in 15 paired nonallergic (NonA) donors from the allergen-response study of Kearns et al. ^18^ Top, median donor-paired differences in relative usage for all eight programmes and three contrasts; the box marks P3. Bottom, P3 paired differences for each donor (connected points, same donor; black bars, medians). Two-sided Wilcoxon signed-rank tests with Benjamini–Hochberg correction across all 24 tests; \**q <* 0.05, \*\**q <* 0.01, \*\*\**q <* 0.001. n.e., not estimable (P10 usage is zero in all donors). There is no unstimulated reference. **d,** Fraction meeting the Gray reconstruction-error criterion (*≤* 0.924). Diamonds denote all-record fractions and circles known donors pooled across samples; labels give record and donor counts where available. Gray is the discovery cohort; PANGEA donor identifiers are unavailable. The criterion evaluates reconstruction, not identity accuracy.

To test whether these programmes capture variation in an independent dataset, we projected them onto activated blood *γδ* T cells from Kearns et al., which were stimulated with cockroach extract, mouse extract or the phosphoantigen HDMAPP.^18^ In 15 nonallergic donors with paired measurements, P3 usage was higher after cockroach-extract stimulation than after either mouse-extract or HDMAPP stimulation (adjusted *P <* 0.001 for both; Fig. 4c). P3 was the only programme with this pattern. Programmes derived from a separate reference therefore captured stimulus-associated transcriptional differences.

Because the programme gene weights are fixed, the reference can be projected onto other datasets, including the stimulus-response and cutaneous T-cell lymphoma cohorts.^18,19^ Reconstruction quality varied across datasets and donors (Fig. 4d); per-cell reconstruction measures are therefore provided to guide downstream interpretation.

### cNMF programmes organize a cross-dataset *γδ* T-cell atlas

To place *γδ* T cells from different studies within a common state space, we used the programme reference to assemble 182,638 cells from eight datasets into a *γδ* T-cell atlas (Fig. 5b). Each record retains its evidence category: TCR-supported, source-annotated or model-predicted. A fixed UMAP was defined from a source-balanced subset of 5,735 cells, with identity scores, TCR subtype, tissue and disease annotations available as overlays (Extended Data Fig. 6).

**Figure 5:**
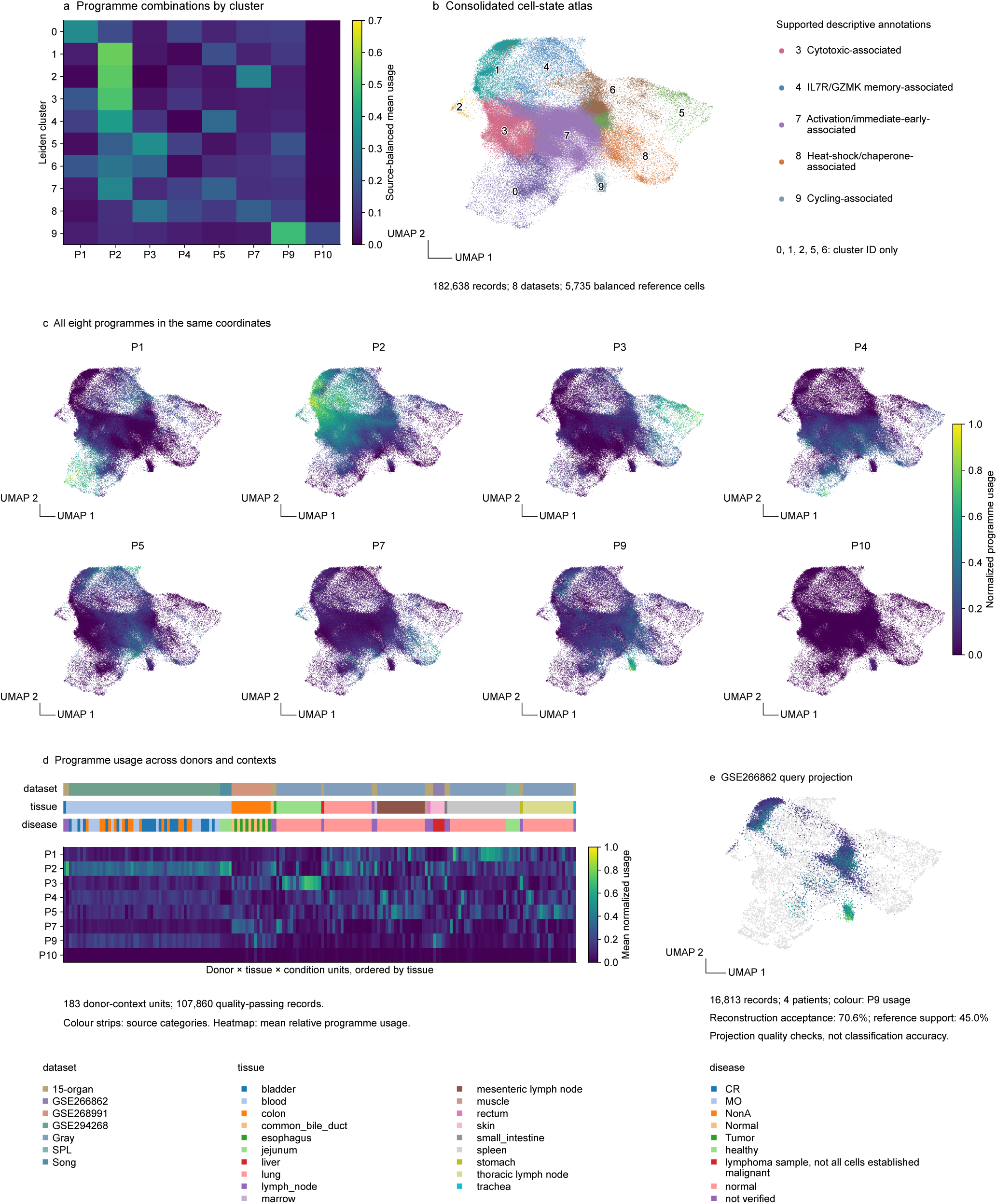
A consolidated programme-space atlas of human *γδ*-associated cells. **a,** Source-balanced programme usage across ten Leiden clusters, restricted to quality-passing records without linked TCR conflicts. **b,** Fixed UMAP of 182,638 records from eight datasets, constructed from 5,735 balanced reference cells. Cluster IDs are retained; descriptive state annotations are limited to clusters 3, 4, 7, 8 and 9. **c,** All eight programmes (P1, P2, P3, P4, P5, P7, P9 and P10), shown as normalized usage in identical coordinates and on a common 0–1 scale. **d,** Mean usage across 183 dataset/donor/tissue/condition units, comprising 107,860 eligible records. Samples receive equal weight within units; unknown donors are excluded. The dataset, tissue and disease/condition colour strips are categorical; complete keys are included below the heatmap. Original labels and capitalization are retained. Heatmap colours represent mean relative usage. **e,** GSE266862 query mapping without refitting the reference UMAP, coloured by P9 usage; grey denotes reference landmarks. The query comprises 16,813 records from four patients. Reconstruction acceptance is 70.6% using the Gray reference error 95th percentile. Exploratory reference support is 45.0%, requiring nearest-reference distance within its reference 95th percentile and a distance-weighted cluster vote of at least 0.6 among 15 reference neighbours. These are separate projection-quality checks, not classification accuracy. Records include distinct TCR-supported, source-annotated and model-candidate categories. Broader overlays are supplied in Extended Data and Supplementary Information. Cross-donor marker support and dominant-dataset exclusion are shown in Supplementary Figs. S1 and S2. Those analyses assess marker persistence within fixed cluster assignments. The query mapping illustrates reuse of the atlas and is not an external validation.

Clustering in programme space yielded ten groups, five of which were annotated by marker expression: cytotoxic (cluster 3), *IL7R*/*GZMK* memory (cluster 4), activation/immediate-early (cluster 7), heat-shock (cluster 8) and cycling (cluster 9) (Fig. 5b; Extended Data Figs. 7–9). To exclude dataset-specific effects, we repeated the marker analysis after removing each cluster’s dominant dataset. Cytotoxic markers and *IL7R* remained elevated across eleven donors in clusters 3 and 4, respectively, and cycling markers persisted in five donors in cluster 9 (Supplementary Table 3; Supplementary Figs. S1 and S2).

Programme usage varied continuously within and across clusters (Fig. 5c,d). To test whether this variation was related to TCR subtype, we compared V*δ*1 and V*δ*2 cells within the same donors. Usage of the immediate-early programme P5 was lower in V*δ*1 than in V*δ*2 cells in all eight eligible donors across three cohorts, whereas P2 and P3 differences were inconsistent in direction (Supplementary Table 4). One lymphoma donor excluded by the reconstruction-quality filter showed the opposite P5 difference; this pattern should therefore not be interpreted as a universal subtype difference.

We next placed the 25 TCR-supported *γδ* T cells recovered from NK-labelled populations (Fig. 3h) on the atlas. All passed the reconstruction criterion. Compared with subtype-matched paired-TCR reference cells from the same donors, these cells showed higher usage of the cytotoxic programme P2 in both HC1 and HC2 (Supplementary Table 5). *γδ* T cells annotated as NK cells were therefore associated with a cytotoxic programme, although this observation is limited to these two donors.

Finally, to demonstrate reuse of the atlas, we mapped *γδ* T cells from a cutaneous T-cell lymphoma study onto the fixed embedding (Fig. 5e).^19^ The atlas provides shared coordinates, programme usage, metadata and projection-quality measures for comparing existing datasets and mapping new ones.

### gdTProfiler identifies candidate *γδ* T-cell-associated bins in spatial transcriptomics data

To test whether gdTProfiler could be applied beyond dissociated cells, we scored all 8-*µ*m bins in the Visium HD colorectal cancer dataset of Faria de Oliveira et al. ^20^ The model identified 39 candidate bins across three tumour sections (Fig. 6a–c). Most candidates (34 of 39) contained detectable *TRDC* RNA (Fig. 6a–f). Because *TRDC* is a model input, this overlap supports, but does not independently validate, the predictions.

**Figure 6:**
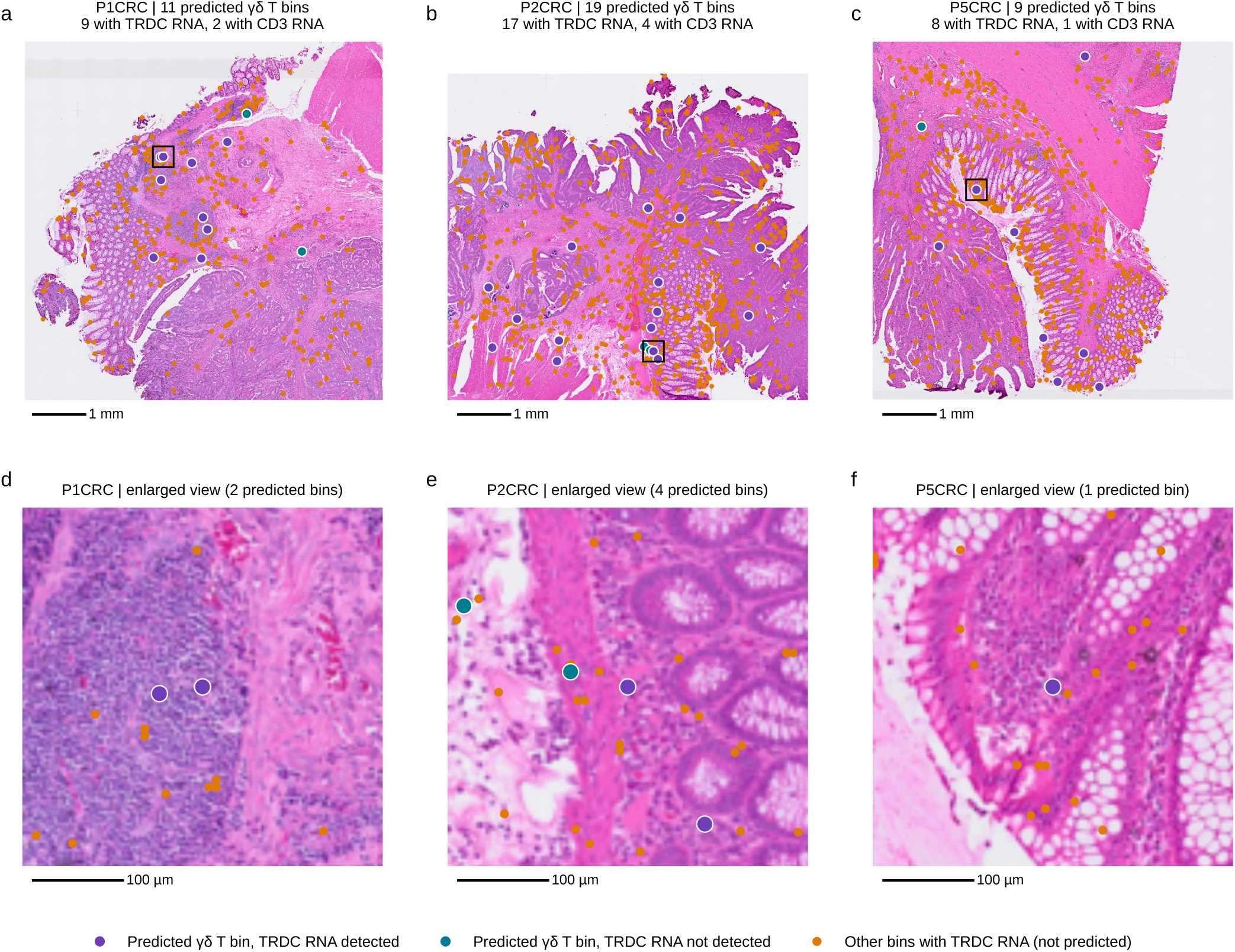
Spatial localization and RNA support of candidate *γδ*-associated bins in colorectal cancer. Visium HD sections P1CRC, P2CRC and P5CRC from the colorectal cancer dataset of Faria de Oliveira et al. ^20^, shown on the original H&E images. **a–c,** All 8-*µ*m bins predicted as *γδ* T cells by gdTProfiler (*p*(*γδ*) *≥* 0.5), scored without annotation-based pre-selection. Purple, predicted bins with detected *TRDC* RNA; teal, predicted bins without detected *TRDC* RNA; orange points, all other bins with detected *TRDC* RNA (371, 882 and 701 bins in P1CRC, P2CRC and P5CRC). Panel titles give the number of predicted bins and how many contain *TRDC* or *CD3D* /*E* /*G* RNA. Black squares mark the regions enlarged in d–f. **d–f,** Enlarged 400 *×* 400-*µ*m views, each centred on the predicted bin with the most other predicted bins nearby; regions were selected without reference to RNA. Detection requires a raw count *>* 0. Scale bars, 1 mm (a–c) and 100 *µ*m (d–f). Bins are spatial measurements, not verified individual cells. *TRDC* is a model input and therefore provides supporting rather than independent evidence; *TRDV1/2/3* were not measured. Prediction–RNA overlays and local views are shown in Extended Data Fig. 10.

Spatial bins can contain RNA from more than one cell, so these candidates mark regions for further investigation rather than verified *γδ* T cells. The programme reference also reconstructed candidate bins poorly, and none met the reference criterion. Candidates in P2CRC showed a trend towards local aggregation (*P* = 0.01, *q* = 0.06), whereas those in P1CRC and P5CRC did not (Supplementary Table 6). Candidates were not closer to cancer-associated fibroblasts than matched T-cell bins (Extended Data Fig. 10).

## Discussion

In this study, we developed gdTProfiler, a hierarchical model that predicts *γδ* T-cell identity and TCR subtype and describes cell state using a fixed programme reference. Previous work identified *γδ* T cells from TCR gene expression; ^8^ gdTProfiler additionally distinguishes *γδ* T cells from NK cells and predicts V*δ*1/V*δ*2 subtype from non-TCR genes. The model generalized across held-out donors and external cohorts and did not classify held-out NK cells as *γδ* T cells. However, 27 TCR-defined *γδ* T cells in SPL were classified as NK cells, and subtype performance varied across cohorts. The programme reference, derived from a separate dataset, captured stimulus-associated variation and organized a cross-dataset atlas.

Our analysis of NK-labelled populations suggests that some *γδ* T cells in public atlases are annotated as NK cells. In HC1 and HC2, all 25 NK-labelled cells with *γδ* TCR sequences were predicted as *γδ* T cells, and these cells showed higher cytotoxic programme usage than other *γδ* T cells from the same donors. This is consistent with the shared cytotoxic features of *γδ* T and NK cells, ^4,5^ which may make such cells difficult to annotate from the transcriptome alone. Candidates without TCR evidence, or with discordant TCR evidence, require confirmation by TCR sequencing or protein measurements.

Unlike discrete cluster labels, continuous programmes allow activities such as cytotoxicity, activation and cycling to coexist within the same cell. ^13^ Because the programme gene weights and embedding coordinates are fixed, new datasets can be projected without refitting the reference. Marker expression was consistent across donors for several clusters, but this does not establish that these clusters represent discrete cell types. In addition, immediate-early and heat-shock genes can be induced during tissue dissociation,^21^ so P5 and P7 usage should not be interpreted as disease-specific activity without further evidence.

Reconstruction quality was lower in the stimulated and lymphoma cohorts than in the discovery reference (Supplementary Table 2).^18,19^ This may reflect expression activities absent from the reference, as well as differences in sequencing depth and sample processing, which our analysis cannot separate. Per-cell reconstruction measures allow poorly represented cells to be identified before state analysis. Deriving programmes from multiple cohorts may improve coverage, provided that the improvement is confirmed in held-out cohorts.

gdTProfiler also extended to spatial transcriptomics, where it identified candidate *γδ* T-cell-associated bins in colorectal cancer sections. Because 8-*µ*m bins can contain RNA from more than one cell and *TRDC* is a model input, these candidates prioritize regions for follow-up rather than confirm *γδ* T-cell identity. Combining gdTProfiler with spatial TCR sequencing or protein imaging would allow these predictions to be validated in tissue.

This study still has some limitations. The external cohorts were examined during model development and therefore provide retrospective tests; evaluation in newly generated datasets will further establish generalization. In addition, because the NK references and T cells came from different studies, NK specificity should be confirmed in datasets that profile both populations together. In future work, we will apply gdTProfiler to additional spatial transcriptomics datasets to investigate the biological significance of *γδ* T-cell localization and state within tissues.

Overall, gdTProfiler provides a common framework for identifying *γδ* T cells, comparing their transcriptional states across datasets and prioritizing cells for TCR-based validation.

## Methods

### Study design and datasets

All analyses were performed on publicly available human single-cell and spatial transcriptomic datasets (Supplementary Table 1), for which ethical approval and informed consent were obtained by the original studies.

The identity model was trained on cells and TCR labels from Song et al., ^8^ together with NK references from GSE139249, ^14^ which were also used for patient-held-out NK evaluation. Subtype labels were taken from Song et al. and GSE268991. ^22^ Spleen cells (SPL) from the Cross-tissue Immune Cell Atlas ^10^ and healthy blood donors HC1 and HC2 from GSE149356^15^ served as external test cohorts and contributed no training labels. NK-labelled populations were analysed in the 15-organ atlas^16^ and PANGEA. ^17^ Programmes were derived from adult *γδ* T cells in Gray et al.^4^ and projected onto Song, SPL, 15-organ, PANGEA, GSE268991, GSE294268 (stimulus response) ^18^ and GSE266862 (cutaneous T-cell lymphoma).^19^ Spatial analyses used Visium HD sections P1CRC, P2CRC and P5CRC from GSE280318. ^20^ Source annotations and repeated biopsies were retained as provided, and cell counts refer to the analysed subsets.

### TCR references and RNA quality control

RNA and TCR records were matched by dataset, donor and normalized cell barcode. Paired *γδ* references were defined as cells with eligible TRG and TRD chains and no TRA or TRB chain, and paired *αβ* references as cells with TRA and TRB chains and no TRG or TRD chain; cells with mixed-lineage chains were excluded from evaluation. In Song et al., eligible chains were those flagged as cell-associated, high-confidence and productive, with non-empty sequences and positive read and UMI counts; full-length status was not required. Contig-level quality fields missing from the processed SPL and GSE149356 releases were not inferred.

Song cells with an eligible TRD chain but no TRG, TRA or TRB chain (TRD-only; *n* = 1,654) were used as training positives but not as evaluation references (Fig. 1a). TRG-only and mixed-chain cells received no *γδ* training label. Subtype labels were assigned from unambiguous TRDV1 or TRDV2 usage; dual or unresolved assignments were excluded. The training pool comprised 23,914 Song and GSE268991 cells, 848 GSE139249 NK references and 3,827 subtype labels, of which 1,639 came from TRD-only cells (34 V*δ*1 and 1,605 V*δ*2). Class imbalance was addressed with sample weights; evaluation prevalence was not altered.

Song and SPL cells were retained if they had at least 200 detected genes, no more than 20% mitochondrial counts, expression of at least one of *CD3D*, *CD3E*, *CD247*, *LCK* or *PTPRC*, and no source doublet flag. This filter removes non-immune and low-quality cells and does not discriminate T cells from NK cells. TCR-reference analysis in SPL was restricted to donors A29, A31, A35, A36 and A37, which had documented TCR assay coverage. Other datasets were used with their original preprocessing and quality control.

### Model architecture and training

The scVI encoder ^23,24^ was trained on quality-controlled Song T-lineage cells using 2,000 variable non-TCR genes, two hidden layers of 128 units, a 20-dimensional latent space and a negative-binomial likelihood with gene-specific dispersion. GSE268991 cells did not contribute to encoder training. Genes beginning with TRA, TRB, TRG or TRD were removed from the variable-gene set; this rule also removed a small number of non-TCR genes. Full gene lists are provided with the code.

The latent mean of each cell was standardized and passed to a shared 20 *→* 20 layer with ReLU activation. Two heads followed. The identity head received the 20 shared features together with nine TCR features (*TRAC*, *TRBC1*, *TRBC2*, *TRDC*, *TRGC1*, *TRGC2*, *TRDV1*, *TRDV2* and *TRDV3*) and produced softmax scores for *αβ* T, *γδ* T and NK cells (29 *→* 3). The subtype head received only the 20 shared features and produced a sigmoid score for V*δ*1 versus V*δ*2 (20 *→* 1). TCR features were normalized to 10^4^ counts within the model gene panel, log-transformed with log(1 + *x*) and standardized with training parameters. Genes absent from a query assay were set to zero and flagged as unmeasured.

The shared layer and subtype head were initialized from a binary *γδ*-versus-*αβ* model trained with the same encoder. Its identity weights initialized the *γδ* row of the three-class softmax layer; the *αβ* row was initialized to zero, and the NK row to zero weights with bias *−*5. With the encoder and standardization fixed, the shared layer and both heads were trained jointly for 100 epochs with Adam^25^ (learning rate 0.001, batch size 256, seed 17), minimizing

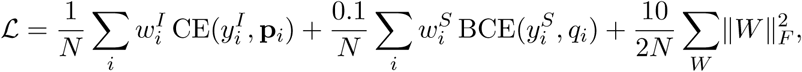

where CE is categorical cross-entropy over the three softmax scores **p***_i_* and BCE is binary cross-entropy for the subtype score *q_i_*. Identity weights 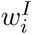 were applied only to Song T cells and GSE139249 NK references and balanced classes, and donors within each class. Subtype weights 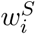 were applied only to cells with eligible subtype labels and balanced donors, and subtypes within each donor. Each weight vector was scaled to mean one over the training table, and biases were not penalized.

### Donor-held-out evaluation and comparators

Identity prediction was evaluated by holding out each of the four Song donors in turn. In each fold, the gene set, encoder, standardization and initialization were derived without the held-out donor, and the model was trained on the remaining Song cells, GSE268991 subtype labels and GSE139249 NK references. Subtype prediction was evaluated over 11 held-out donors (four Song and seven GSE268991). All cells from a held-out donor were excluded from training. NK specificity was assessed in five GSE139249 patient-held-out fits. ^14,26^ No external data were used to select parameters or thresholds.

The model trained on all source donors was applied without refitting to SPL and HC1/HC2 (GSM4498553 and GSM4498554). Because HC1/HC2 contain only *γδ* references, identity performance in these cohorts is reported as sensitivity.

Comparators were logistic regression (scikit-learn defaults,^27^ with TCR genes excluded for subtype prediction), CellTypist (Immune_All_Low) ^10^ and an adaptation of the modular TCR score of Song et al.; ^8^ implementation details are given in Supplementary Methods.

AUROC and AP were calculated with *γδ* T cells (identity) and V*δ*1 cells (subtype) as positive classes; subtype metrics were calculated within TCR-defined V*δ*1/V*δ*2 cells. *γδ* calls used a threshold of 0.5 on the softmax score. Because donors are the unit of biological replication, ^28^ donor-level metrics are also reported. Training variability was assessed by retraining with five seeds (17–21).

### Analysis of NK-labelled populations

The trained model from Fig. 2 was applied without refitting to the 15-organ and PANGEA datasets, using a threshold of 0.5 and sensitivity thresholds of 0.7 and 0.8. Original annotations and UMAP coordinates were retained, and cells without coordinates were included in all counts. The 15-organ close-up panels show the region containing the author-labelled T GD Cell, NK/T Cell and T/NK clusters (UMAP1 1.5–16.8; UMAP2 *−*0.21–10.21). PANGEA includes GSE139249 records that were used in training. For HC1/HC2, TCR support required matched TRG/TRD and CDR3 records in the PANGEA-processed data, and these analyses used the PANGEA expression objects rather than the GEO files used in Fig. 2.

Marker expression was normalized to 10^4^ counts per cell and log-transformed. Means were calculated within organs (15-organ) or study–sample units (PANGEA) and averaged equally across units. Because some markers are also model inputs, their expression provides supporting rather than independent evidence.

### Programme discovery and projection

Consensus NMF (cNMF) ^13^ was applied to a donor-balanced set of 5,077 adult *γδ* T cells from 22 donors in Gray et al., with 200 NMF runs per tested rank, and *K* = 8 was selected (Extended Data Fig. 5). Programmes were numbered according to their matches in an initial *K* = 10 solution (cosine similarity *≥* 0.976), giving P1–P5, P7, P9 and P10; the *K* = 10 programmes P6 and P8 were absorbed into P1 (cosine 0.96) and P7 (0.84), respectively.

For projection, counts were normalized within the reference gene set to 10^6^ and divided by reference gene standard deviations, without centring or log transformation. With the programme-by-gene spectra *H* fixed, non-negative usage coefficients were estimated for each cell *x_i_*,

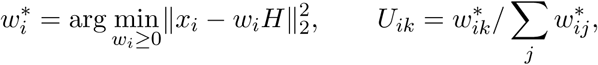

and cells with zero total usage were left undefined. Projection required at least 80% coverage of the 2,000 reference genes. A cell passed the reconstruction criterion if its relative reconstruction error did not exceed the 95th percentile of the Gray reference (0.924).

Hallmark overlap ^29^ was tested using the top 50 genes per programme, with Benjamini–Hochberg correction over 800 tests. ^30^ In GSE294268, donor-paired programme usage was compared between stimuli (cockroach extract, mouse extract and HDMAPP; 15 nonallergic donors) using two-sided Wilcoxon signed-rank tests, ^31^ with Benjamini–Hochberg correction over 24 programme–contrast tests. The study did not include an unstimulated condition.

### Atlas construction and clustering

The atlas combined eight datasets and retained source identifiers, donor and sample metadata, TCR evidence and prediction scores for each record. To reduce duplication, PANGEA studies already included as separate datasets (GSE149356, GSE268991, GSE294268, GSE266862, GSE263248 and GSE164378) were removed from the PANGEA contribution. Low-quality or uncertain records were flagged rather than removed.

The UMAP ^32^ was fitted to square-root-transformed programme usage of 5,735 reference cells, sampled with at most 150 cells per donor or PANGEA study (Euclidean distance, 30 neighbours, minimum distance 0.3, seed 17). Reference cells required an appropriate source annotation, adequate reconstruction and no conflicting TCR record; GSE294268 and GSE266862 were mapped as queries. Leiden clustering^33^ was performed in the eight-programme space (resolution 0.5, seed 17), and remaining records were assigned by weighted voting among their 15 nearest reference neighbours. Seeds 18 and 19 and resolutions 0.25 and 1 were used as sensitivity analyses. Assignments were considered well supported if the neighbour vote was at least 0.6 and the distance to the nearest reference cell was within the reference 95th percentile.

Within-sample marker contrasts required at least ten cells per group, used log-normalized counts and were averaged within donors and then equally across donors and datasets. To test whether cluster markers depended on a single dataset, the dataset contributing most records to each cluster was removed and marker summaries were recomputed with clusters and UMAP unchanged. In Fig. 5d, 107,860 records from 183 donor-context units were weighted equally within dataset, donor, tissue and condition; records without a donor identifier were excluded.

Subtype comparisons used cells with unambiguous paired-TCR V*δ*1/V*δ*2 labels and adequate reconstruction, excluding conflicting TCR records. Each sample required at least ten cells of each subtype. Differences in mean usage (V*δ*1 minus V*δ*2) were averaged across samples within donors and then across donors within cohorts. Sensitivity analyses removed the reconstruction criterion or added TRD-only references. Wilcoxon signed-rank tests with Benjamini–Hochberg correction were applied to cohorts with at least five donors; cohorts with two to four donors are reported descriptively.

HC1/HC2 cells were projected onto the same programme reference and UMAP without refitting, separately from the atlas in Fig. 5b. The 25 NK-labelled cells with matched TRG/TRD and CDR3 records formed the TCR-supported group. Within each donor, the remaining paired-TCR cells provided subtype-specific reference means, weighted by the subtype composition of the TCR-supported group. Comparisons in the 15-organ and PANGEA datasets used the original NK/T, T/NK and NK annotations within matched organ or study–sample units.

### Spatial analysis

The model scored all 1,595,565 filtered 8-*µ*m bins in P1CRC, P2CRC and P5CRC without pre-selection by annotation; 1,511,849 bins had sufficient counts for scoring. The primary threshold was *p*(*γδ*) *≥* 0.5, with 0.7 and 0.8 as sensitivity thresholds. RNA detection was defined as a positive raw count, and H&E images and Space Ranger coordinates were used unchanged. Close-up examples were the bins closest to the median candidate score, selected without reference to RNA evidence. *TRDV1–3* were not measured in these sections, so subtype predictions were not evaluated.

Local aggregation was measured as the fraction of candidate bins with another candidate within 100 *µ*m and compared with 199 permutations that preserved candidate counts within 500-*µ*m blocks and RNA-depth quartiles, with Benjamini–Hochberg correction across comparisons; sections with fewer than two candidates were not tested. Candidate bins were projected onto the fixed *K* = 8 programme reference and assessed with the Gray reconstruction criterion. In P2CRC, candidates were matched to source-annotated CD4/CD8 T-cell bins by tissue region, spatial block, RNA depth and distance to the tumour boundary, and their proximity to source-annotated cancer-associated fibroblast (CAF) bins was compared (Supplementary Methods).

### Software and reproducibility

Analyses were performed in Python 3.13 using scvi-tools 1.5.2, PyTorch 2.12, Scanpy 1.12, scikit-learn 1.9, SciPy 1.17, pandas 2.3, umap-learn 0.5 and leidenalg 0.12, ^24,27,32–36^ together with cNMF 1.7.1^13^ and CellTypist 1.7. ^10^ Module scores for the Song et al. comparator were computed in R 4.6 with Seurat 5.5. ^12^ The trained model parameters, input gene lists and the *K* = 8 programme reference are distributed with the gdTProfiler package, together with a tutorial notebook and the software requirements (see Code availability). Analysis scripts used to generate the figures are available from the corresponding authors on reasonable request.

## Data availability

Song expression and TCR data are available through Zenodo record 7989561. ^8^ SPL data derive from E-MTAB-11536. ^10^ HC1/HC2 and the 15-organ atlas are available under GSE149356 and GSE159929, respectively.^15,16^ NK references are available under GSE139249. ^14^ PANGEA ^17^ is available through https://pangeapy.org/ and Figshare version 2 (https://doi.org/10.6084/m9.figshare.28138364.v2).^37^ Released TCR records were matched at the sample and cell level rather than assumed available for all atlas records.

Gray adult expression data are available in the CELLxGENE collection Human *γδ* T cells in diverse tissues exhibit site-specific maturation dynamics across the lifespan. ^4^ The processed adult object contains 7,417 cells before the analysis-specific discovery selection. GSE240858 contains pediatric expression and GSE263248 tissue TCR repertoire data. Additional source and projection datasets are GSE268991, GSE294268 and GSE266862. ^18,19,22^ CRC spatial data are available under GSE280318. ^20^ GEO accessions can be accessed at https://www.ncbi.nlm.nih.gov/geo/. Programme gene weights and model parameters are included in the software archive (https://doi.org/10.5281/zenodo.24486259).

## Code availability

gdTProfiler is available at https://github.com/KaelynWenHKU/gdTProfiler and v1.0.0 is archived at Zenodo (https://doi.org/10.5281/zenodo.24486259).

### Acknowledgements

We thank Jaeyoung Shim for early-stage exploration of the project and Jong-Eun Park for the generous early sharing of the full PANGEA dataset.

This work was supported by the National Research Foundation of Korea (NRF) grant funded by the Korea government (MSIT) (RS-2023-00239434).

## Author contributions

K.N.W. developed the computational workflow, performed single-cell and spatial transcriptomic analyses, and wrote the original draft. Y.H. led the supervision of the study. S.H.P. proposed the initial idea for the study and co-supervised the work.

